# Thyroid hormone signaling specifies cone subtypes in human retinal organoids

**DOI:** 10.1101/359950

**Authors:** Kiara C. Eldred, Sarah E. Hadyniak, Katarzyna A. Hussey, Boris Brennerman, Pingwu Zhang, Xitiz Chamling, Valentin M. Sluch, Derek S. Welsbie, Samer Hattar, James Taylor, Karl Wahlin, Donald J. Zack, Robert J. Johnston

## Abstract

The mechanisms underlying the specification of diverse neuronal subtypes within the human nervous system are largely unknown. The blue (short-wavelength/S), green (medium-wavelength/M) and red (long-wavelength/L) cone photoreceptors of the human retina enable high-acuity daytime vision and trichromatic color perception. Cone subtypes are specified in a poorly understood two-step process, with a first decision between S and L/M fates, followed by a decision between L and M fates. To determine the mechanism controlling S vs. L/M fates, we studied the differentiation of human retinal organoids. We found that human organoids and retinas have similar distributions, gene expression profiles, and morphologies of cone subtypes. We found that S cones are specified first, followed by L/M cones, and that thyroid hormone signaling is necessary and sufficient for this temporal switch. Temporally dynamic expression of thyroid hormone degrading and activating proteins supports a model in which the retina itself controls thyroid hormone levels, ensuring low signaling early to specify S cones and high signaling late to produce L/M cones. This work establishes organoids as a model for determining the mechanisms of cell fate specification during human development.

**One sentence summary:** Cone specification in human organoids

---

Cone photoreceptors in the human retina enable daytime, color, and high acuity vision (*1*). The three subtypes of human cones are defined by the visual pigment that they express: blue- (short-wavelength/S), green- (medium-wavelength/M), or red- (long-wavelength/L) opsin (*2*). Specification of human cones occurs in a two-step process. First, a decision occurs between S vs. L/M cone fates (**Fig. 1A**). If the L/M fate is chosen, a subsequent choice is made between expression of L- or M-opsins (*3-6*). Mutations affecting opsin expression or function cause various forms of color blindness and retinal degeneration (*7-9*). Great progress has been made in our understanding of the vertebrate eye through the study of model organisms. However, little is known about the developmental mechanisms that generate the mosaic of mutually exclusive cone subtypes in the human retina. Here, we study the specification of human cone subtypes using human retinal organoids differentiated from stem cells (**Fig. 1D-K**).

**Figure 1.**
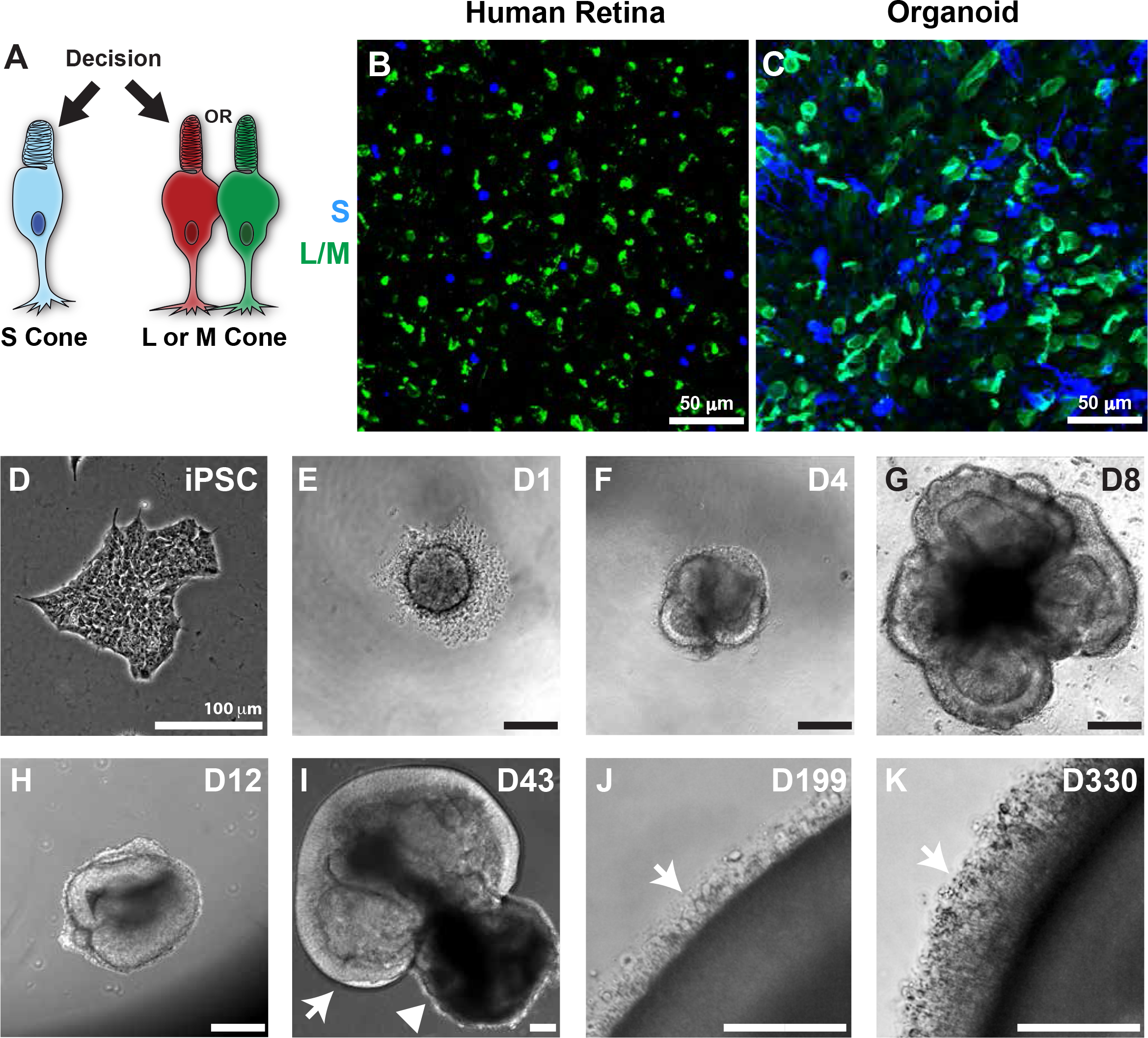
S and L/M cone generation in human retinal organoids. **A)** Decision between S and L/M cone subtype fate. **B-C)** S-opsin (blue) L/M-opsin (green). **B)** Human adult retina age 53. **C)** iPSC-derived organoid, day 200 of differentiation. **D-K)** Bright field images of organoids derived from iPSCs. **D)** Undifferentiated iPSCs. **E)** Day 1: aggregation. **F)** Day 4: formation of neuronal vesicles. **G)** Day 8: differentiation of retinal vesicles. **H)** Day 12: manual isolation of retinal organoid. **I)** Day 43: arrow indicates developing retinal tissue, arrowhead indicates developing retinal pigment epithelium (RPE). **J)** Day 199: arrow indicates outer segments. **K)** Day 330: arrow indicates outer segments.

Human retinal organoids generate photoreceptors that respond to light (*10-14*). We find that human organoids recapitulate the specification of cone subtypes observed in the human retina, including the temporal generation of S cones followed by L and M cones. Moreover, we find that this regulation is controlled by thyroid hormone signaling, which is necessary and sufficient to control cone subtype fates through the nuclear hormone receptor Thyroid Hormone Receptor β (Thrβ). Expression of thyroid hormone-regulating genes suggests that retina-intrinsic temporal control of thyroid hormone levels and activity governs cone subtype specification. While retinal organoids have largely been studied for their promise of therapeutic applications (*15*), our work demonstrates that human organoids can also be used to reveal fundamental mechanisms of human development.

## Specification of cone cells in organoids recapitulates development in the human retina

We compared features of cone subtypes in human organoids to adult retinal tissue. Adult human retinas and organoids at day 200 of differentiation displayed similar ratios of S to L/M cones as indicated by expression of S- or L/M-opsins (adult: S=13%, L/M=87%; organoid: S=29%, L/M=71%)(**Fig. 1B-C, S1A**). We examined L/M cones with an antibody that recognizes both L- and Mopsin proteins due to their extremely high similarity. Both S and L/M cones expressed the cone-rod-homeobox transcription factor (CRX), a critical transcription factor for photoreceptor differentiation (**Fig. 2A, E**)(*16-18*), indicating proper fate specification in organoids. Additionally, cones in organoids and retinas displayed similar morphologies, with L/M cones that had longer outer segments and wider inner segments than S cones (**Fig. 2B-D, F-H**)(*19*). The outer segments of cones were shorter in organoids than in adult retinas, consistent with postnatal maturation (**Fig. 2D, H**)(*20*). Thus, cone subtypes in human retinal organoids displayed distributions, gene expression patterns, and morphologies similar to cones of the human retina.

**Figure 2.**
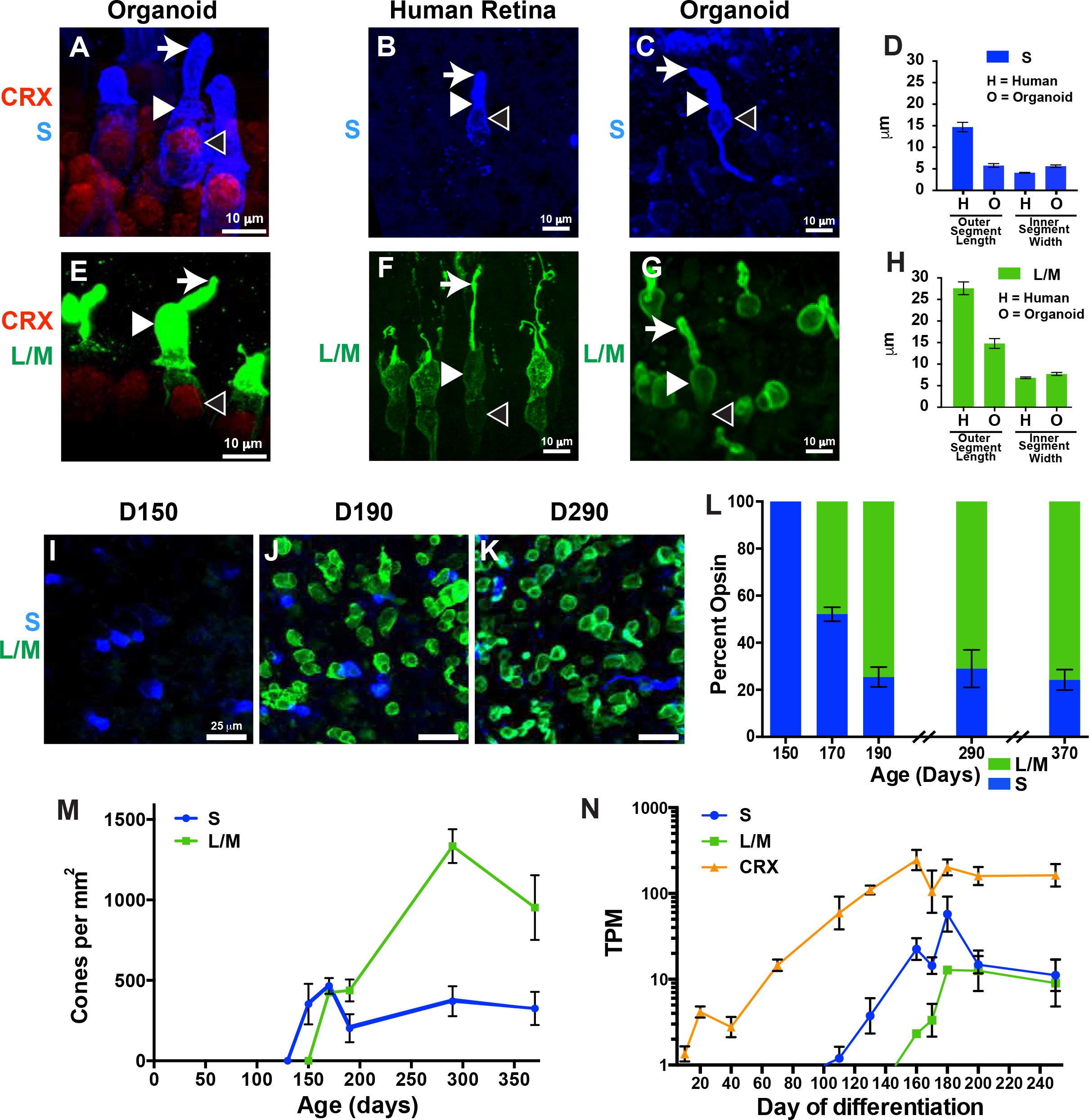
Human cone subtype specification is recapitulated in organoids. **A-K)** S-opsin (blue) and L/M-opsin (green) were examined in human iPSC-derived organoids (**2A, C-E, G-M**) and human retinas (**2B, D, F, H**). **A-C, E-G)** Arrows indicate outer segments, full arrowheads indicate inner segments, empty arrowheads indicate nuclei. **A,E)** CRX (a general marker of photoreceptors) is expressed in S cones and L/M cones. **B-D)** S cones display short outer segments and thin inner segments in both human retinas and organoids. **F-H)** L/M cones display long outer segments and wide inner segments in both human retinas and organoids. **D,H)** Quantification of outer segment lengths and inner segment widths (adult retina: L/M, n=13, S, n=10; organoid: L/M, n=35, S, n=42). **I-N)** S cones are generated before L/M cones in organoids. **L)** Ratio of S:L/M cones during organoid development. **M)** Density of S and L/M cones during organoid development. **N)** *S-opsin* expression precedes *L/M-opsin* expression in human iPSC-derived organoids. *CRX* expression starts before opsin expression. TPM=Transcripts per Kilobase Million.

We next examined the developmental dynamics of cone subtype specification in organoids. In the human retina, S cones are generated during fetal weeks 11-34 (days 77-238), whereas LM cones are specified later during fetal weeks 14-37 (days 98-259)(*21, 22*). We tracked the ratios and densities of S and L/M cones in organoids by antibody staining over 360 days of differentiation. A significant number of cones expressing S-opsin were first observed at day 150 (**Fig. 2I, L-M**). The density of S cones leveled off at day 170 (**Fig. 2M**), at the timepoint when cones expressing L/M-opsin began to be observed (**Fig. 2J-M**). The population of L/M cones increased dramatically until day 300 (**Fig. 2K-M**) when they reached a steady-state density. Remarkably, the 20-day difference between S- and L/M-opsin expression onset in retinal organoids is similar to the 20-day difference observed in the appearance of S- and L/M- cones in the fetal retina (*21*). These observations show a temporal switch from S cone specification to L/M cone specification during retinal development.

We next conducted RNA-Seq through 250 days of iPSC-derived organoid development. We found that *S-opsin* RNA was expressed first at day 111 and leveled off at day 160, while *L/M-opsin* RNA was expressed at day 160 and remained steady after day 180, consistent with the timeline of photoreceptor maturation in organoids and fetal retinas (**Fig. 2N, Fig S1B**). Moreover, *CRX* RNA and CRX protein were expressed before opsins in organoids, similar to human development (*23*) (**Fig. 2N, Fig. S1B-G**,). Thus, human organoids recapitulate many aspects of the developmental timeline of cone subtype specification observed in human retinas, providing a model system to uncover the mechanisms of these developmental changes.

## Thyroid hormone signaling is necessary and sufficient for the temporal switch between S and L/M fate specification

Seminal work in mice identified thyroid hormone receptor β2 (Thrβ2) as a critical regulator of cone subtype specification: *Thrβ2* mutants display a complete loss of M-opsin expression and a complete gain of S-opsin expression in cone photoreceptors (*24-26*). Similar roles for *Thrβ2* have been characterized in other organisms with highly divergent cone patterning (*27-29*). Additionally, rare human mutations in *Thrβ2* are reported to alter color perception, indicative of a change in the S to L/M cone ratio (*30*). To directly test the role of *Thrβ2* in human cone subtype specification, we used CRISPR/Cas9 in human embryonic stem cells (ESCs) to generate a homozygous mutation resulting in early translational termination in the unique first exon of *Thrβ2* (**Fig. S2A**). Surprisingly, organoids derived from these mutant stem cells displayed no changes in cone subtype ratio or density (wild-type: S=62%, L/M=38%; *Thrβ2* KO: S=59%, L/M=41%; P=0.83), indicating that, unlike previous suggestions based on other species, *Thrβ2* is dispensable for cone subtype specification in humans (**Fig. 3A-C**).

**Figure 3.**
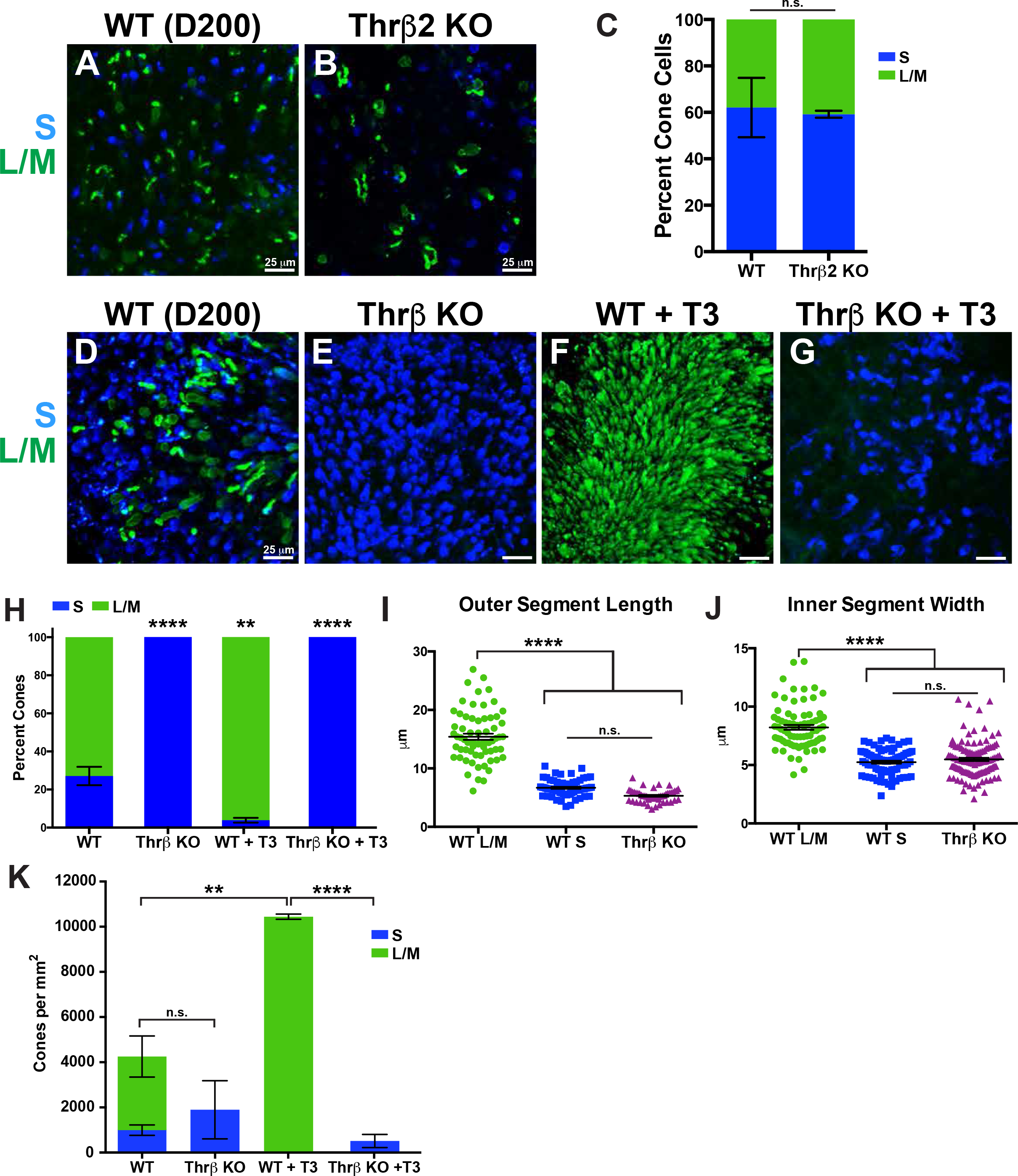
Thyroid hormone signaling is necessary and sufficient for the temporal switch between S and L/M fate specification. **A-K)** S-opsin (blue) and L/M-opsin (green) were examined in human ESC-derived organoids. **A)** Wild-type (WT) **B)** *Thrβ2* early termination mutant (*Thrβ2 KO*). **C)** Quantification of A-B (WT n=3, *Thrβ2 KO* n=3) **D)** Wild-type (WT) **E)** *Thrβ* Knockout (*Thrβ* KO) **F)** WT treated with 20 nM T3 (WT + T3). **G)** *Thrβ* KO treated with 20 nM T3 (*Thrβ* KO + T3). **H)** Quantification of D-E (WT, n=9; *Thrβ* KO, n=3; WT + T3, n=6; *Thrβ* KO + T3, n=3. Tukey’s multiple comparisons test comparisons test: WT vs Thrβ KO, P<0.0001; WT vs WT + T3, P < 0.01; WT + T3 vs Thrβ KO + T3, P<0.0001). **I)** Length of outer segments (WT, L/M n=66 cells, WT, S n=66 cells, *Thrβ* KO, n=50 cells. Tukey’s multiple comparisons test: WT L/M vs. WT SW, P<0.0001; WT L/M vs. *Thrβ* KO, P<0.0001; WT S vs. *Thrβ* KO, not significantly different). **J)** Width of inner segments (WT, L/M n=78 cells; WT, S n=78 cells; *Thrβ* KO, n=118 cells. Tukey’s multiple comparisons test: WT L/M vs. WT SW, P<0.0001; WT L/M vs. *Thrβ* KO, P<0.0001; WT S vs. *Thrβ* KO, not significantly different). **K)** T3 acts through Thrβ to increase total cone number. Quantification of density of S and L/M cones. (WT, n=6; *Thrβ* KO, n=3; WT + T3, n=3; *Thrβ* KO + T3, n=3. Tukey multiple comparisons test between total cone numbers: WT vs. *Thrβ* KO, not significantly different; WT vs WT + T3, P<0.01; WT + T3 vs *Thrβ* KO + T3, P<0.0001).

Since *Thrβ2* alone is not required for human cone subtype specification, we reexamined data from Weiss *et. al* (*30*) and found that missense mutations in exons 9 and 10 affected both *Thrβ2* and another isoform of the human *Thrβ* gene, *Thrβ1* (**Fig. S2A**). Thus, we asked whether *Thrβ1* and *Thrβ2* together are required for cone subtype specification in humans. To completely ablate Thrβ function (i.e. Thrβ1 and Thrβ2), we used CRISPR/Cas9 in human ESCs to delete a shared exon that codes for part of the DNA-binding domain (DBD) of *Thrβ* (**Fig. S2A**). *Thrβ* null mutant retinal organoids displayed a complete conversion of all cones to the S subtype (wild-type: S=27%, L/M=73%; *Thrβ* KO: S=100%, L/M=0%; P<0.0001) (**Fig. 3D-E, H**). In these mutants, all cones expressed S-opsin and had the S cone morphology (**Fig. 3I-J**). Thus, Thrβ is required to activate L/M and to repress S cone fates in the human retina.

Thrβ binds with high affinity to triiodothyronine (T3), the more active form of thyroid hormone, to regulate gene expression (*31*). Since L/M cones differentiate after S cones, we hypothesized that T3 acts through Thrβ late in retinal development to induce L/M cone fate and repress S cone fate. One prediction of this hypothesis is that addition of T3 early in development will induce L/M fate and repress S fate. To test this model, we added 20nM T3 to ESC- and iPSC-derived organoids starting from days 20-50 to day 200 of differentiation. We observed a dramatic conversion of cone cells to L/M fate (wild-type: S=27%, L/M=73%; wild-type + T3: S=4%, L/M=96%; P<0.01) (**Fig. 3F, H, Fig. S2B**). Thus, early addition of T3 is sufficient to induce L/M fate and suppress S fate.

To test whether T3 acts specifically through Thrβ to control cone subtype specification, we differentiated *Thrβ* mutant organoids with early T3 addition. *Thrβ* mutation completely suppressed the effects of T3, generating organoids with only S cones (wild-type + T3: S=4%, L/M=96%; *Thrβ* KO + T3: S=100%, L/M=0%; P<0.0001) (**Fig. 3F-H**). We conclude that T3 acts though Thrβ to promote L/M cone fate and suppress S cone fate.

We confirmed the regulation of L/M-opsin expression through thyroid hormone signaling in a retinoblastoma cell line, which expresses L/M-opsin when treated with T3 (**Fig. S2C-D**)(*32*). T3-induced activation of *L/M-opsin* expression was suppressed upon RNAi knock down of *Thrβ* (**Fig. S2E-F**), similar to the suppression observed in human organoids.

Interestingly, in organoids, early T3 addition not only converted cone cells to L/M fate but also dramatically increased cone density (**Fig. 3F, K**). Moreover, T3 acts specifically through Thrβ to control cone density (**Fig. 3G, K**). Early T3 addition may increase cone density by advancing and extending the temporal window of L/M cone generation.

Together, these results demonstrate that T3 signals though Thrβ to promote L/M cone fate and repress S cone fate in developing human retinal tissue.

## Dynamic expression of thyroid hormone-regulating genes during development

Our data suggest that temporal control of thyroid hormone signaling determines the S vs. L/M cone fate decision, whereby low signaling early induces S fate and high signaling late induces L/M fate. Thyroid hormone exists largely in two states: Thyroxine (T4), the most abundant circulating form of thyroid hormone, and T3, which binds thyroid hormone receptors with high affinity (*31, 33*). Conversion of T4 to T3 occurs locally in target tissues to induce gene expression responses (*34, 35*). Deiodinases, enzymes that modulate the levels of T3 and T4, are expressed in the retinas of mice, fish, and chicken (*29, 36-40*). Therefore, we predicted that T3- and T4-degrading enzymes would be expressed during early human eye development to reduce thyroid hormone signaling and specify S cones, while T3-producing enzymes, carriers, and transporters would be expressed later in human eye development to increase signaling and generate L/M cones.

To test these predictions, we examined gene expression across 250 days of organoid development. The expression patterns of thyroid hormone-regulating genes were grouped into three classes: changing expression (**Fig. 4A**), consistent expression (**Fig. 4B**), or no expression (**Fig. 4C**). Interestingly, Deiodinase 3 (*DIO3*), an enzyme that degrades T3 and T4 (*34*), was expressed at high levels early in organoid development but at low levels later (**Fig. 4A**). Conversely, Deiodinase 2 (*DIO2*), an enzyme that converts T4 to active T3 (*34*), was expressed at low levels early but then dramatically increased over time (**Fig. 4A**). We examined RNA-Seq data from Hoshino *et. al* (*23*) and found that developing human retinas display similar temporal changes in expression of *DIO3* and *DIO2* (**Fig. S3A**). Deiodinase 1 (*DIO1*), which regulates T3 and T4 predominantly in the liver and kidney (*41*), was not expressed in organoids or retinas (**Fig. 4C, S3C**). Thus, the dynamic expression of *Dio3* and *Dio2* supports low thyroid hormone signaling early in development to generate S cones and high thyroid hormone signaling late to produce L/M cones.

**Figure 4.**
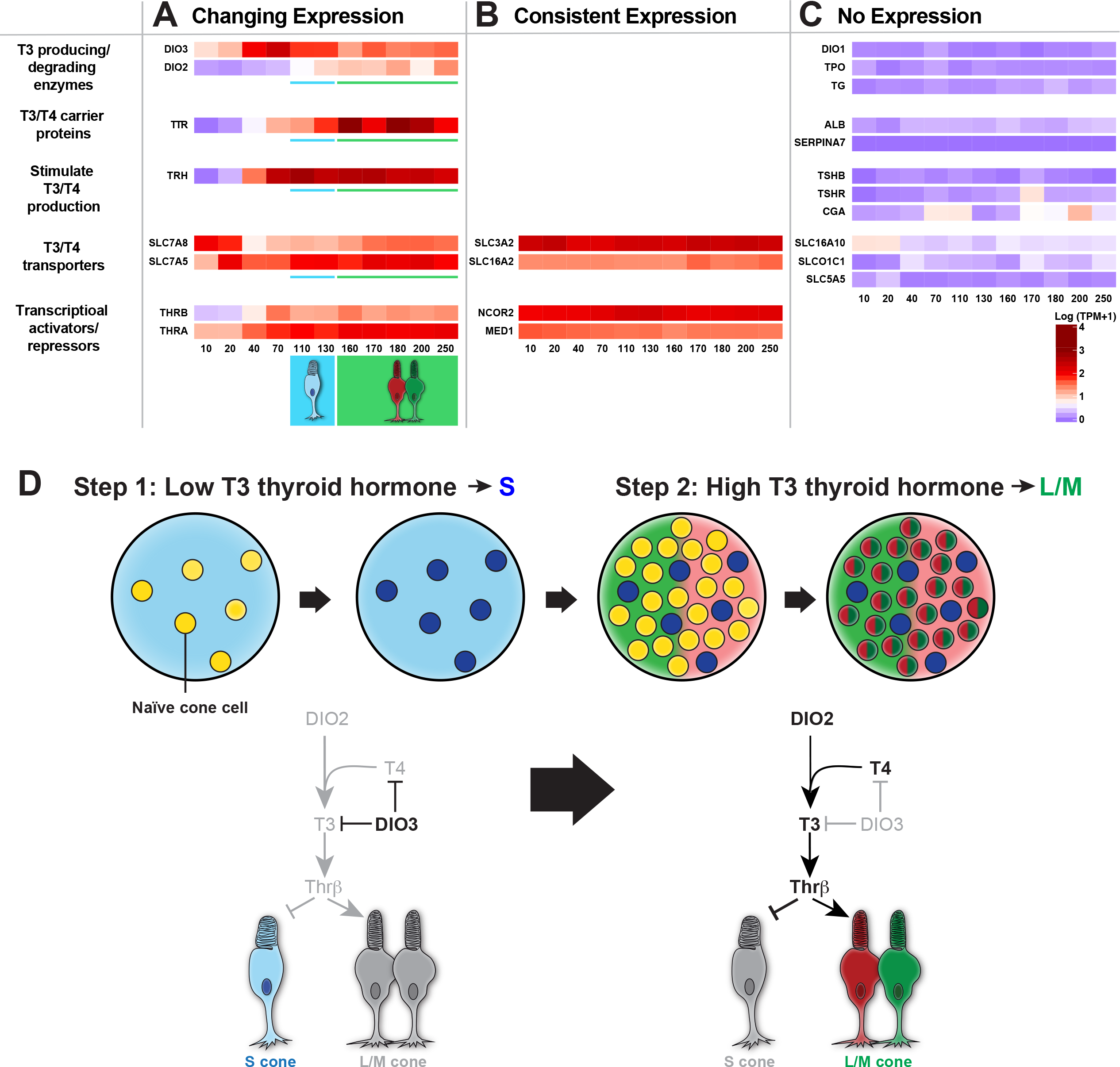
Dynamic expression of thyroid hormone signaling regulators during development. **A-C)** Heat maps of Log(Transcripts per Kilobase Million (TPM) + 1) values for genes with (A) changing expression, (B) consistent expression, and (C) no expression. Numbers at the bottom of heat maps indicate organoid age in days. **D)** Model of the temporal mechanism of cone subtype specification in humans. For simplicity, only the roles of DIO3 and DIO2 are illustrated. In step 1, expression of DIO3 degrades T3 and T4 leading to S cone specification. In step 2, expression of DIO2 converts T4 to T3 to signal Thrβ to repress S and induce L/M cone fate.

Consistent with a role for high thyroid hormone signaling in the generation of L/M cones later in development, expression of transthyretin (*TTR*), a thyroid hormone carrier protein, increased during organoid and retinal development (**Fig. 4A, S3A**)(*23*). In contrast, albumin (*ALB*) and thyroxine-binding globulin (*SERPINA7*), other carrier proteins of T3 and T4, were not expressed in organoids or retinas (**Fig. 4C, S3C**)(*23*).

T3 and T4 are transported into cells via membrane transport proteins (*42*). The T3/T4 transporters *SLC7A5* and *SLC7A8* increased in expression during organoid differentiation (**Fig. 4A**). Additionally, two T3/T4 transporters, *SLC3A2* and *SLC16A2*, were expressed at high and consistent levels throughout organoid development (**Fig. 4B**). Other T3/T4 transporters (*SLC16A10, SLCO1C1, SLC5A5*) were not expressed in organoids (**Fig. 4C**), suggesting tissue-specific regulation of T3/T4 uptake. We observed similar expression patterns of T3/T4 transporters in human retinas (**Fig. S3A-C**)(*23*).

We next examined expression of transcriptional activators and repressors that mediate the response to thyroid hormone. Consistent with *Thrβ* expression in human cones (*43*), expression of *Thrβ* in organoids increased with time as cone cells were specified (**Fig. 4A**). Expression of thyroid hormone receptor a (*Thrα*) similarly increased with time (**Fig. 4A**). Thyroid hormone receptor cofactors, co-repressor *NCoR2* and co-activator *MED1*, were expressed at steady levels during organoid differentiation (**Fig. 4B**). Similar temporal expression patterns were observed in human retinas (**Fig. S3A-B**)(*23*). Thus, our data suggest that expression of Thrβ and other transcriptional regulators enables gene regulatory responses to differential thyroid hormone levels.

A complex pathway controls production of thyroid hormone. Thyrotropin-releasing hormone (TRH) is produced by the hypothalamus and other neural tissue. TRH stimulates release of thyroid-stimulating hormone *α* (CGA) and thyroid-stimulating hormone β (TSHβ) from the pituitary gland. CGA and TSHβ bind the thyroid-stimulating hormone receptor (TSHR) in the thyroid gland. T3 and T4 production requires Thyroglobulin (TG), the substrate for T3/T4 synthesis, and Thyroid Peroxidase (TPO), an enzyme which iodinates tyrosine residues in TG (*44*). Interestingly, *TRH* was expressed in organoids and retinas but the other players were not (**Fig. 4A-C, S3A-C**)(*23, 45, 46*), suggesting that the retina itself does not generate thyroid hormone, rather it modulates the relative levels of T3 and T4 and expresses TRH to signal for thyroid hormone production in other tissues.

Therefore, the temporal expression of thyroid hormone signaling regulators supports our model that the retina intrinsically controls T3 and T4 levels, ensuring low thyroid hormone signaling early to promote S fate and high thyroid hormone signaling late to specify L/M fate (**Fig. 4D**).

Organoids provide a powerful system to determine the mechanisms of human development. Model organism and epidemiological studies generate important hypotheses about human biology that are often experimentally intractable. This work shows that organoids enable direct testing of hypotheses in developing human tissue.

Our studies identify temporal regulation of thyroid hormone signaling as a mechanism controlling cone subtype specification in humans. Consistent with our findings, preterm human infants with low T3/T4 have an increased incidence of color vision defects (*47-50*). Moreover, our identification of a mechanism that generates one cone subtype while suppressing the other is critical for developing organoid-based therapies to treat diseases such as color blindness, retinitis pigmentosa, and macular degeneration.

